# GDV1 C-terminal truncation of 39 amino acids disrupts sexual commitment in *Plasmodium falciparum*

**DOI:** 10.1101/2020.10.28.360123

**Authors:** Marta Tibúrcio, Eva Hitz, Igor Niederwieser, Gavin Kelly, Heledd Davies, Christian Doerig, Oliver Billker, Till S. Voss, Moritz Treeck

## Abstract

Malaria is a mosquito-borne disease caused by apicomplexan parasites of the genus *Plasmodium.* Completion of the parasite’s life cycle depends on the transmission of sexual stages, the gametocytes, from an infected human host to the mosquito vector. Sexual commitment occurs in only a small fraction of asexual blood stage parasites and is initiated by external cues. The gametocyte development protein 1 (GDV1) has been described as a key facilitator to trigger sexual commitment. GDV1 interacts with the silencing factor heterochromatin protein 1 (HP1), leading to its dissociation from heterochromatic DNA at the genomic locus encoding AP2-G, the master transcription factor of gametocytogenesis. How this process is regulated is not known. In this study we have addressed the role of protein kinases implicated in gametocyte development. From a pool of available protein kinase KO lines, we identified two kinase knockout lines which fail to produce gametocytes. However, independent genetic verification revealed that both kinases are not required for gametocytogenesis but both lines harbour the same mutation that leads to a truncation in the extreme C-terminus of GDV1. Introduction of the identified nonsense mutation into the genome of wild type parasite lines replicates the observed phenotype. Using a GDV1 overexpression line we show that the truncation in the GDV1 C-terminus does neither interfere with the nuclear import of GDV1 nor its interaction with HP1 *in vitro*, but appears important to sustain GDV1 protein levels and thereby sexual commitment.

**Importance:** Transmission of malaria causing *Plasmodium* species by mosquitos requires the parasite to change from a continuously growing asexual parasite form growing in the blood, to a sexually differentiated form, the gametocyte. Only a small subset of asexual parasites differentiates into gametocytes that are taken up by the mosquito. Transmission represents a bottleneck in the lifecycle of the parasite, so a molecular understanding of the events that lead to stage conversion may identify novel intervention points. Here we screened a subset of kinases we hypothesized to play a role in this process. While we did not identify kinases required for sexual conversion, we identified a mutation in the C-terminus of the Gametocyte Development 1 protein (GDV1), which abrogates sexual development. The mutation destabilises the protein but not its interaction with its cognate binding partner HP1. This suggest an important role for the GDV1 C-terminus beyond trafficking and protein stability.

## Introduction

Malaria is a devastating disease caused by parasites of the genus *Plasmodium*, leading to ~ 405,000 deaths per year ^1^. *Plasmodium falciparum* causes the most severe and life-threatening form of human malaria. The complex life cycle involves interactions with multiple tissues in two different organisms, the human host and the mosquito vector. Inside the human host *P. falciparum* predominantly infects red blood cells (RBC) where it asexually replicates or, a small fraction (0-20%), commits to sexual development (gametocytogenesis) ^2^. Gametocytogenesis occurs preferentially in the extravascular compartment in the bone marrow and spleen ^3–9^. After 10-12 days mature stage V gametocytes are released into the peripheral circulation to allow transmission to mosquitoes.

Sexual commitment can be initiated by metabolic cues in the human host. Specifically, it has been described that depletion of lysophosphatidylcholine (LysoPC), a common component of human serum, leads to increased rates of gametocyte production and therefore represents the first molecularly defined factor known to inhibit or trigger sexual conversion ^10^. Sexual commitment depends on upregulation of the *ap2-g* gene ^2,11^, which requires removal of heterochromatin protein 1 (HP1) from chromatin. HP1 interacts directly with the gametocyte development 1 protein (GDV1), which causes HP1 to dissociate ^12^. HP1 is responsible for repression of a range of genes ^13^, while GDV1 specifically acts on the *ap2-g* locus. How this specificity is achieved is not known. Furthermore, how a drop in LysoPC levels is sensed and transduced into GDV1-mediated HP1 removal is not understood.

Kinases are key transducers of signals in cellular processes in various stages of the *Plasmodium* life cycle ^14,15^ and are likely candidates to play important roles in gametocyte commitment and development. A study by Solyakov et al ^14^ has identified a panel of likely and confirmed non-essential protein kinases, some of which are transcribed during sexual development (PlasmoDB) or in gametocytes^16–19^. Aiming to identify protein kinases involved in sexual development we screened eight KO lines for phenotypes in gametocyte induction and/or maturation. Two lines made no gametocytes, but subsequent validation showed that their gametocytogenesis defect was not due to the absence of these kinases. Instead, we found that both lines shared the same truncation in the C-terminal end of GDV1, which caused the loss of gametocyte development. Here, we address the importance and role of the GDV1 C-terminal for sexual commitment and interaction with HP1. We show that the loss of the C-terminal 39 amino acids of GDV1 does not interfere with nuclear import and interaction with HP1 in vitro, but prevents GDV1 from triggering efficient sexual commitment.

## Results

### Identification and characterization of *Plasmodium falciparum* kinase KO lines with a gametocytogenesis phenotype

It has been shown previously in *P. berghei* that protein kinases non-essential during the asexual blood stages are essential in other lifecycle stages, for example during parasite transmission in the mosquito ^15^. To identify kinases important for gametocytogenesis we investigated the role of a group of likely non-essential kinases ^14^ during asexual blood stages development. Using the lines described by Solyakov *et al ^14^,* which have been generated by single crossover gene disruption, we induced sexual development using conditioned medium ^20^ and followed progression through the stages I to V of gametocytogenesis (Figure 1a). Six of the eight KO lines displayed normal gametocyte development, while two, TKL2 (PF3D7_1121300) and eIK2 (PF3D7_0107600) kinase KO lines, produced very few (≤ 0.1%) gametocytes (Figure 1b). Of these, one has a disrupted tyrosine kinase like 2 (*tkl2*) locus, which has been characterized as a protein kinase secreted outside of the red blood cell ^17^. Gene loss and accumulation of mutations is frequently observed in parasite lines kept in continuous *in vitro* culture over time and the loss of the ability to form gametocytes is not uncommon ^21^. To exclude mutations in the *ap2-g* gene, which was identified through a loss of function mutant previously ^2^, we sequenced the *ap2-g* locus in the 3D7/TKL2 KO parasite line. The sequencing results confirmed that the phenotype observed was not associated with mutations in *ap2-g*, leading us to conclude that the deletion of TKL2 was possibly the cause for the observed phenotype. In order to verify the role of TKL2 in gametocyte induction, we generated a DiCre-mediated TKL2 conditional KO line in NF54 parasites (NF54/TKL2:loxPint). We used CRISPR/Cas9 to simultaneously introduce a DiCre cassette into the *pfs47* locus, as previously described ^22,23^, and to flank the kinases domain of *tkl2* with two loxPints (Figure 1c, Supplementary Figure 1a and b). To address the role of TKL2 in gametocyte development we treated the NF54/TKL2:loxPint line with DMSO (control) or rapamycin (KO) (Supplementary Figure 1b). We then induced sexual commitment using conditioned medium ^20^, and monitored gametocyte development. No difference in commitment or development between the control and the rapamycin-induced NF54/TKL2:loxPint parasites (Figure 1d) were observed. These results show that TKL2 is not involved in sexual commitment or gametocyte development/maturation and another mutation is likely the cause for the observed phenotype.

**Figure 1.**
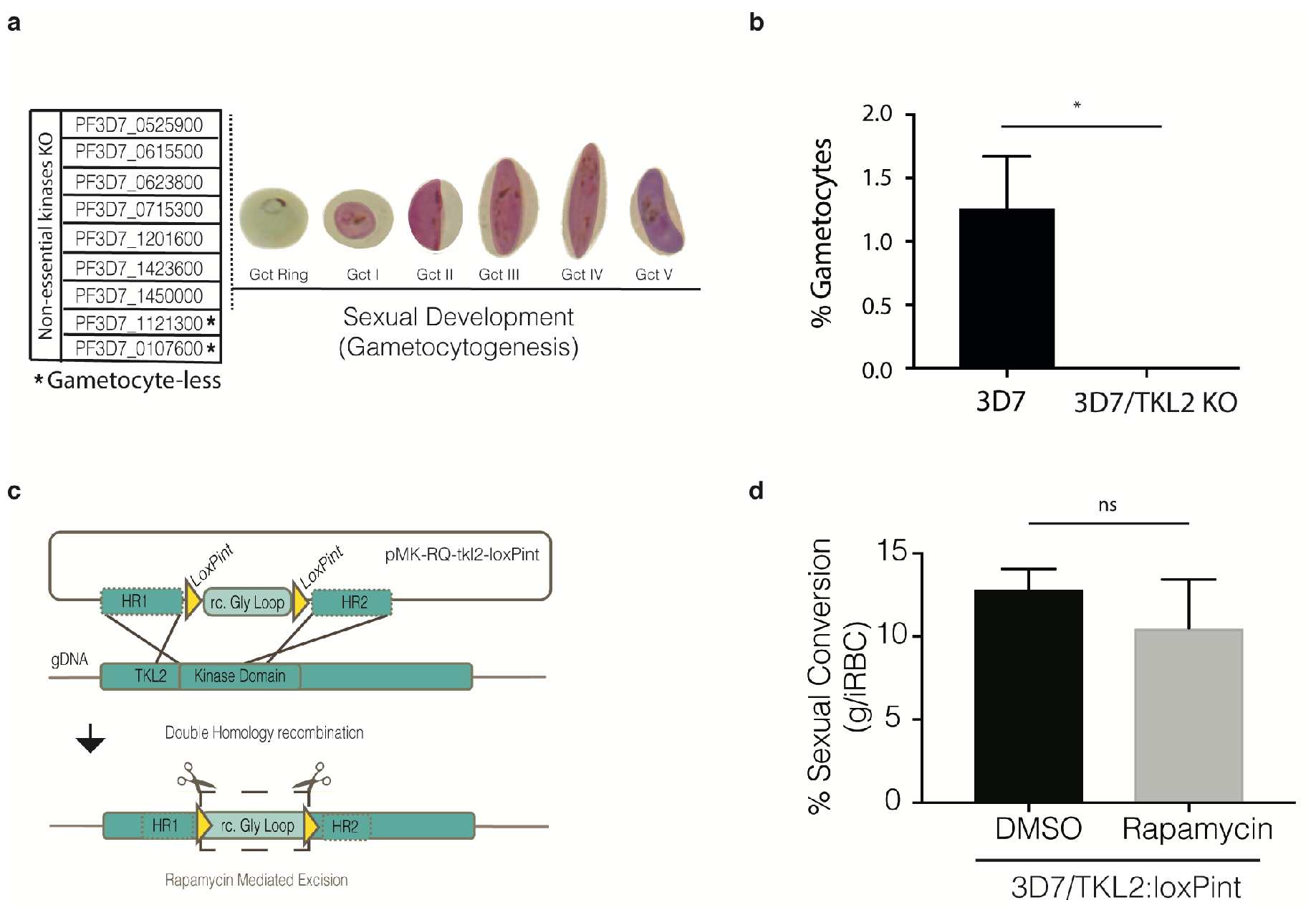
Screening of Plasmodium falciparum non-essential kinases during sexual commitment and development. A) List of non-essential kinases characterized during sexual development in this study (PF3D7_0107600 – eIK2; PF3D7_0525900 - NEK2; PF3D7_0615500 – CRK5; PF3D7_0623800 – TKL4; PF3D7_0715300 - calcium/calmodulin-dependent protein kinase, putative; PF3D7_1121300 – TKL2; PF3D7_1201600 - NEK3; PF3D7_1423600 - calcium-dependent protein kinase, putative; PF3D7_1450000 - serine/threonine protein kinase, putative) ^14^. B) Comparison of the percentage of gametocytaemia between the 3D7 WT line and the PfTKL2 kinase KO clones of the same transfection (clones B10 and B12) generated by single crossover integration ^14^. Each column represents the mean of triplicate microscope counts, each of at least 500 cells, analysed using paired t test, ± SD, (*; p<0.05; 3D7 versus TKL2 KO clones, P=0.0377). C) Schematic of the CRISPR/Cas9 strategy used to generate a TKL2 conditional knockout (KO) line (3D7/TKL2:loxPint) as well as the primers used to confirm successful gene editing (Supplementary Table 2); The *pMK-RQ-tkl2-loxPint* donor plasmid contains a recodonized version of the glycine loop in the kinase domains of *tkl2* (rc. Gly Loop) flanked by two loxPints and homology regions for homology-directed repair. D) Sexual conversion rates in 3D7/TKL2:loxPint parasites treated with DMSO (control) or rapamycin (KO). Each column represents the mean of triplicate microscope counts, each of at least 500 cells, analysed using paired t test ± SD, (ns, p≥0.05; 3D7/TKL2:loxPint treated with DMSO versus Rapamycin, P-0.4017).

### A common GDV1 truncation is found in both kinase KO lines deficient in gametocyte formation

The second kinase KO where a gametocytogenesis defect was identified was the eukaryotic initiation factor serine/threonine kinase 2 (eIK2) KO line (3D7/eIK2 KO) (Supplementary Figure 2a). eIK2 has previously been characterized as non-essential during sexual development in *P. falciparum* and *P. berghei* and elK2 KO lines appeared to undergo normal gametocyte development in rodent *Plasmodium* species ^24^. This indicated that, as 3D7/TKL2 KO, the 3D7/eIK2 KO line also harbours a mutation preventing efficient gametocyte development. Sequencing of the *ap2-g* locus in this parasite line as described above showed no mutations in the coding region of *ap2-g*. Therefore, a potentially unknown mutation underlies the loss of gametocytes in these parasite lines.

To understand the nature of the block in sexual development we analysed the transcriptome of induced wildtype 3D7 parasites and in two eIK2 KO clones using RNAseq. Samples were collected for RNA extraction between 28-32 hours post-invasion (hpi) after induction with conditioned medium (Figure 2a). The RNAseq analysis revealed a significant downregulation in 3D7/eIK2 KO parasites of genes known to be upregulated during gametocytogenesis, including genes that have been shown to be AP2-G-dependent ^2,10,12,25–27^ (Figure 2b and Supplementary Table 1). We found *ap2-g* itself to be downregulated in 3D7/eIK2 KO parasites, but this reached significance only in one of the clones. Together with the lack of mutations in *ap2-g* itself, these results suggested that the block in gametocytogenesis was upstream of AP2-G function during sexual commitment. At that time, GDV1 was shown to be an upstream activator of AP2-G expression ^12^ so we sequenced the *gdv1* locus in the elK KO clones and identified a nonsense mutation in *gdv1* that results in a premature stop codon leading to a C-terminal truncation of 39 amino acids (GDV1Δ39) (Figure 2c and Supplementary Figure 2b). Sequencing of the 3D7/TKL2 KO parasite clones showed the same mutation (Supplementary Figure 2b), suggesting that the deletion of the last 39 amino acids of GDV1 in both mutant lines is responsible for the gametocytogenesis phenotype observed in both kinase KO lines.

**Figure 2.**
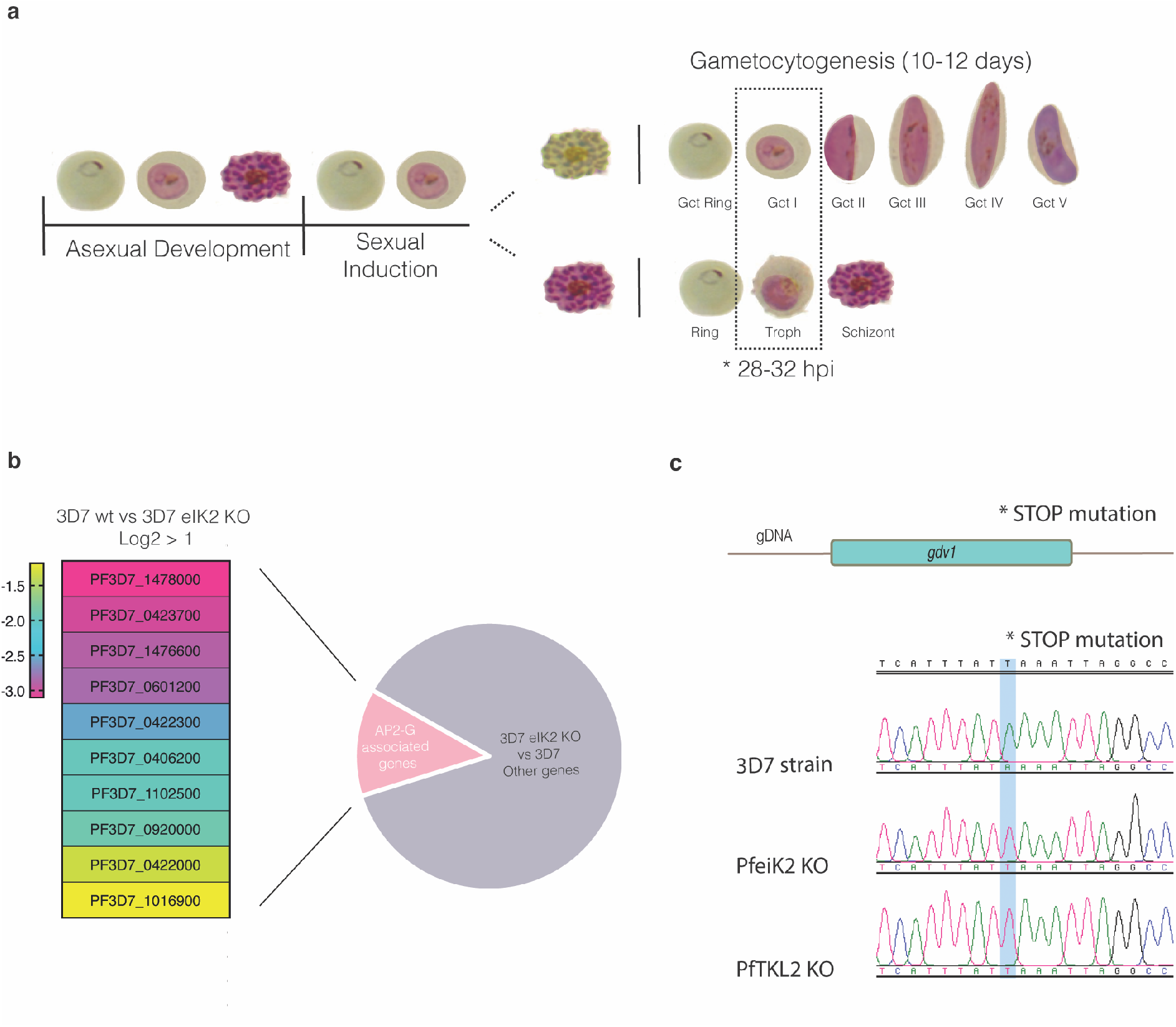
RNA sequencing analysis comparing WT and PfeIK2 gametocyte-less kinase KO line. A) Representation of the interactive cycles of asexual and sexual differentiation upon sexual induction; dotted box illustrates the time point and asexual and sexual stages of the parasite collected for RNAseq. B) Heatmap showing genes previously described as being associated with AP2-G expression and significantly downregulated in the PfeIK2 kinase KO clones (log2 fold change >1). C) DNA sequence trace showing the stop mutation identified in the PfeIK2 and PfTKL2 KO clones which is absent in the 3D7 reference parasite line.

#### The carboxy-terminal 39 amino acids of GDV1 are important for its function

To verify genetically the identified mutation in *gdv1* we generated an 3xHA-tagged version of GDV1Δ39 and introduced it in the endogenous *gdv1* locus in the NF54 parasite line (NF54/GDV1Δ39:HA) (Figure 3a and Supplementary Figure 3a and b). GDV1Δ39:HA parasites lost the ability to form gametocytes (Figure 3b), suggesting the GDV1 C-terminus plays an essential role during sexual commitment or development. Determination of the localisation or expression levels of GDV1Δ39:HA was not possible, as we could not confidently distinguish true signal from background fluorescence. We repeatedly failed to obtain parasites expressing 3xHAtagged full length GDV1 from the endogenous locus to compare expression levels and the localisation of the truncated GDV1 version. Notably, direct C-terminal tagging of GDV1 at the endogenous locus was also not successful in other studies, unless when in combination with a destabilisation domain ^12,28^.

**Figure 3.**
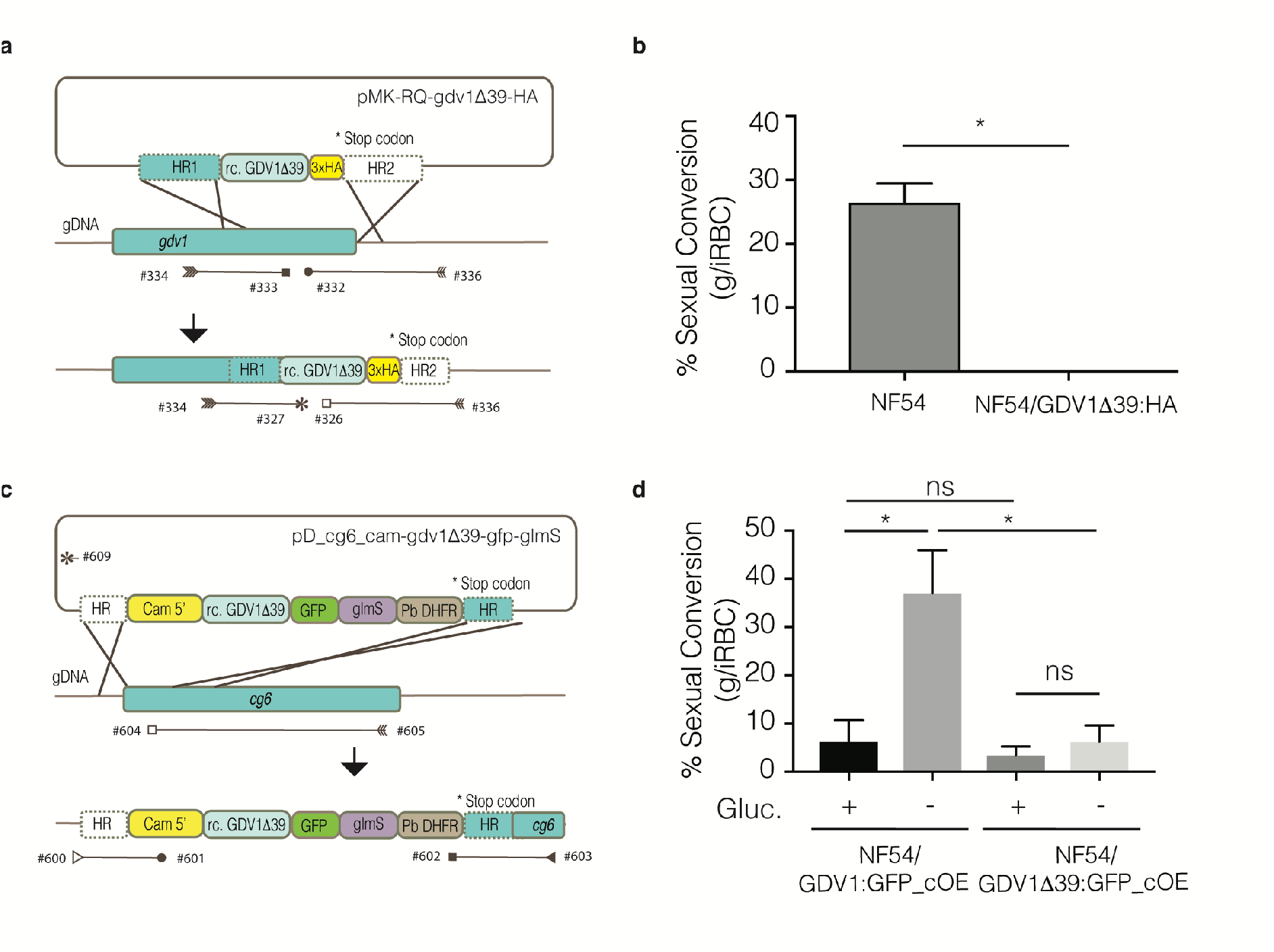
Quantification of gametocyte production in GDV1β39 mutant parasite lines. A) Illustration of the strategy used to generate the 3xHA-tagged GDV1Δ39 mutant line (NF54/GDV1Δ39:HA) as well as the primers used to confirm integration (Supplementary Table 2); The *pMK-RQ-gdv1Δ39-HA* donor plasmid contains a recodonized version of the *gdv1Δ39* mutant and a 3xHA tag, flanked by homology regions. B) Comparison of sexual conversion rates between NF54 and NF54/GDV1Δ39:HA parasite lines. Each column represents the mean of duplicate (NF54) and triplicate (NF54/GDV1Δ39:HA) microscope counts, each of at least 500 cells, analysed using paired t test, ± SD, (*, p<0.05; NF54 versus NF54/GDV1Δ39:HA, P=0.0489). C) Schematic of the strategy used to make the NF54/GDV1Δ39:GFP_cOE overexpressing line as well as the primers used to verify integration of the transgene cassette into the *cg6* (*glp3*) locus (Supplementary Table 2). The *pD_cg6_cam-gdv1Δ39-gfp-glmS* donor plasmid contains a recodonized version of the *gdv1Δ39* mutant followed by the in-frame *gfp* sequence and the *glmS* ribozyme element, flanked by homology regions. D) Comparison of sexual conversion rates between NF54/GDV1:GFP_cOE and NF54/GDV1Δ39:GFP_cOE parasite lines in the presence (prevents sexual conversion) or absence of glucosamine (induces sexual conversion). Each column represents the mean of triplicate counts of at least 500 cells, analysed using paired t test, ± SD, (*, p<0.05; ns, not significant, p≥0.05; NF54/GDV1:GFP_cOE non-induced versus induced, P=0.0094; non-induced NF54/GDV1:GFP_cOE versus NF54/GDV1Δ39:GFP_cOE, P=0.2276; NF54/GDV1Δ39:GFP_cOE non-induced versus induced, P=0.4038; induced NF54/GDV1:GFP_cOE versus NF54/GDV1Δ39:GFP_cOE, P=0.0281).

Therefore, we resorted to a system that allows robust testing of GDV1-dependent gametocyte induction. We introduced an ectopic *gdv1-gfp* fusion gene under control of the calmodulin promoter and a *glmS* ribozyme in the 3’ untranslated region has been introduced into the *cg6* (*glp3*, PF3D7_0709200) locus, allowing conditional overexpression of GDV1-GFP to trigger high gametocyte conversion rates (NF54/iGP2 line) ^29^. In the presence of glucosamine, the *glmS* ribozyme destabilizes the mRNA preventing GDV1:GFP expression, while in the absence of glucosamine GDV1:GFP is overexpressed, leading to gametocyte induction ^30^. For simplicity, the NF54/iGP2 line described by Boltryk and colleagues ^29^ has been renamed NF54/GDV1:GFP_cOE in this study (cOE stands for conditional over expression). To test GDV1Δ39 function in this assay, we introduced a *gdv1Δ39-gfp-glmS* cassette into the *cg6* locus, generating a conditional GDV1Δ39:GFP overexpression parasite line (NF54/GDV1Δ39:GFP_cOE) (Figure 3c and Supplementary Figure 3c and d). We then compared the sexual conversion rates in the NF54/GDV1:GFP_cOE and NF54/GDV1Δ39:GFP_cOE parasite lines in the presence and absence of glucosamine. Unlike the NF54/GDV1:GFP_cOE control line, the induced overexpression of GDV1Δ39:GFP in the NF54/GDV1Δ39:GFP_cOE parasite line failed to trigger a significant increase in sexual commitment (Figure 3d). These results suggest that the full integrity of the GDV1 C-terminus is important for sexual commitment.

#### GDV1Δ39 is imported into the nucleus and retains the ability to interact with HP1

GDV1 is a nuclear protein and we hypothesized that the deletion of a predicted C-terminal nuclear bipartite localisation sequence (cNLS mapper, http://nls-mapper.iab.keio.ac.jp/cgi-bin/NLS_Mapper_form.cgi) may interfere with GDV1 nuclear localisation and hence its ability to interact with HP1 at heterchromatic loci. Therefore we localised GDV1Δ39:GFP in the NF54/GDV1Δ39:GFP_cOE parasites by immunofluorescence at 28-32 hpi (Figure 4a and Supplementary Figure 3e). The results show a clear punctate and nuclear signal in the induced NF54/GDV1:GFP_cOE parasite line, as previously reported (Supplementary Figure 3e) ^12^. NF54/GDV1Δ39:GFP_cOE parasites also show a localized GFP signal in the nucleus, but the signal is weaker and more diffuse when compared with NF54/GDV1:GFP_cOE (Supplementary Figure 3e). In order to quantify and compare GFP levels in NF54/GDV1:GFP_cOE and NF54/GDV1Δ39:GFP_cOE parasites, we performed a whole cell protein extraction for Western blot (WB) analysis using anti-GFP using anti-HSP70 antibodies as a loading control (Figure 4b and c). The WB showed a clear reduction of GDV1Δ39:GFP levels compared to GDV1:GFP (Figure 4c). To quantify the localization of GDV1Δ39:GFP in the cytoplasm compared to the nucleus we prepared cytosolic and nuclear protein extracts using subcellular fractionation (Figure 4b and d). We determined the cytoplasmic fraction using anti-aldolase antibodies ^31^ and anti-histone 3 antibodies were used to determine the nuclear fraction ^32^. GDV1Δ39:GFP was only detected in the nuclear fraction further supporting that its nuclear localisation is not affected by the C-terminal truncation (Figure 4d). Thus, GDV1Δ39:GFP protein level is much reduced compared to GDV1:GFP, despite being expressed from the same locus and driven by the same promoter. To test if the GDV1Δ39 deletion affects its interaction with HP1, we performed an in vitro assay where 6xHIS-tagged GDV1 WT and Δ39 versions were co-expressed with Strep-tagged HP1 in *Escherichia coli* bacteria. Interaction between GDV1 and HP1 is detected by affinity purification of HIS:GDV1 and analysis of co-eluted proteins by Coomassie staining ^12^. HIS-tagged SIP2 does not interact with HP1 and was used as a negative control (Figure 4e). As previously shown, HIS-GDV1 pulled down HP1, which was not observed when SIP2 was used as a bait ^12^. Interestingly, the Δ39:GFP of GDV1 also pulled down HP1, indicating GDV1 C-terminus was not essential for the interaction in *E. coli* (Figure 4e). This observation indicates that the interaction of GDV1Δ39:GFP and HP1 can still occur in the parasite, but that it is insufficient to trigger gametocytogenesis. An explanation for this could be that GDV1Δ39:GFP levels do not reach the threshold required for efficient gametocyte induction. To examine expression of the GDV1Δ39 mutant, we analysed the mean fluorescence intensity in uninduced and induced NF54/GDV1:GFP_cOE and NF54/GDV1Δ39:GFP_cOE parasites at the single cell level using flow cytometry (Figure 4f and g). As expected, NF54/GDV1:GFP_cOE parasites show a robust increase of GFP fluorescence upon induction of GDV1:GFP expression through glucosamine removal. A measurable increase of the mean fluorescence was also observed upon induction in most NF54/GDV1Δ39:GFP_cOE parasites, but well below the levels observed for NF54/GDV1:GFP_cOE parasites. However, a small proportion of NF54/GDV1Δ39:GFP_cOE parasites displayed GFP fluorescence at the level observed in the NF54/GDV1:GFP_cOE control line. In line with its nuclear localisation, GDV1Δ39:GFP may contribute to form gametocytes in these parasites. This is further discussed below.

**Figure 4.**
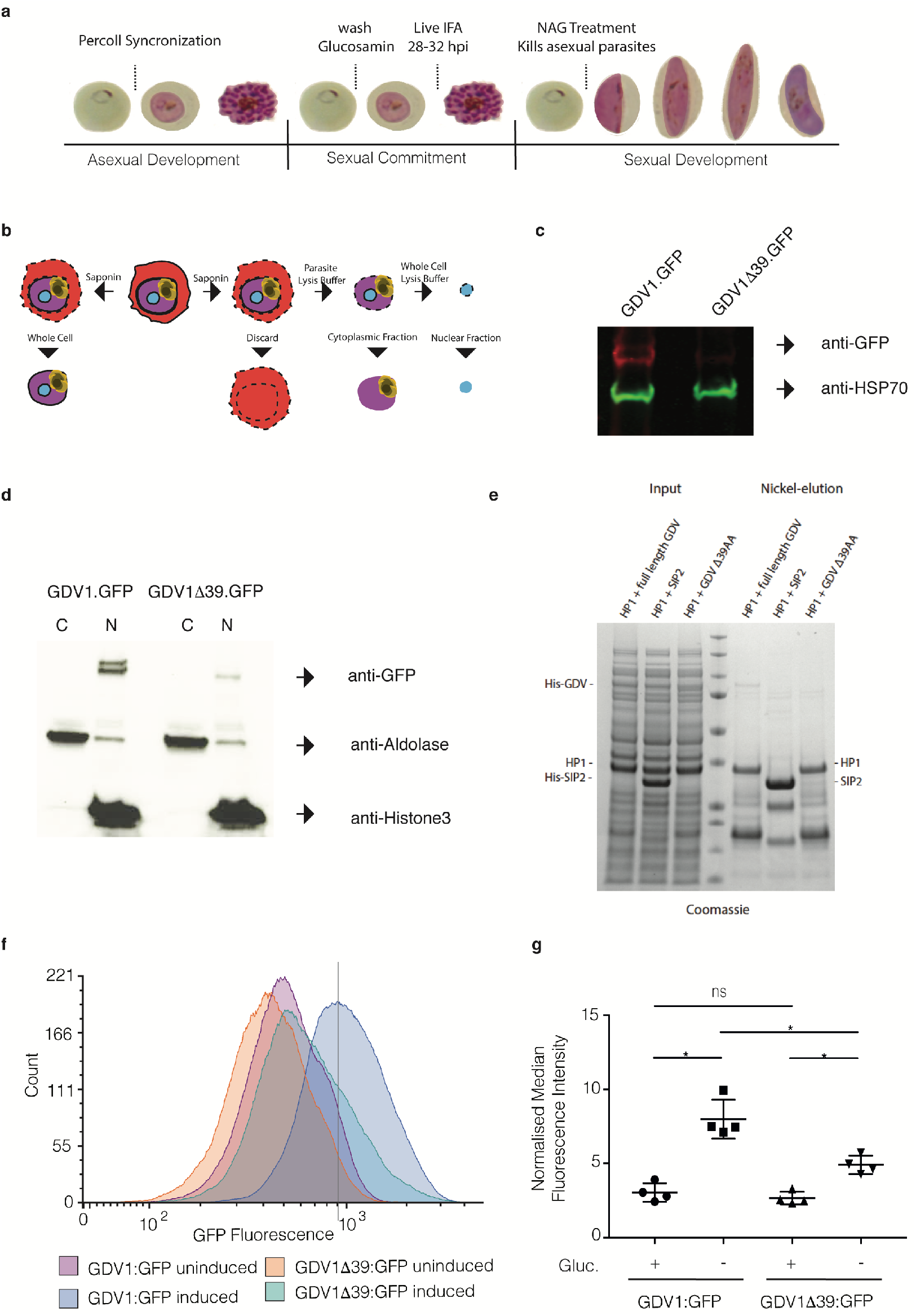
GDV1β39 expression, localization and interaction with HP1. A) Representation of the protocol used to collect the samples used to characterize expression and localization of GDV1Δ39:GFP. B) Illustration of the subcellular fractionation workflow. C) Western blot showing the levels of GDV1:GFP expression in NF54/GDV1:GFP_cOE and NF54/GDV1Δ39:GFP_cOE parasites grown in the absence of glucosamine (induces expression); GDV1:GFP/ GDV1Δ39:GFP expression is detected using an anti-GFP antibody while anti-HSP70 antibodies have been used as controls. D) Western blot showing GDV1:GFP expression levels in the cytoplasmic and nuclear fraction in NF54/GDV1:GFP_cOE and NF54/GDV1Δ39:GFP_cOE parasites cultured in the absence of glucosamine (induces expression). E) Strep-HP1 co-purifies with both HIS-GDV1 and HIS-GDV1Δ39 but not with the HIS-SIP2 control. Coomassie-stained SDS-polyacrylamide gel from pull down experiment with HIS-GDV1/Strep-HP1 and HIS-SIP2/Strep-HP1. Lane 4: protein size standard. F) Representative normalised flow cytometry histograms quantifying GDV1:GFP fluorescence for each parasite line. The experiment was repeated 4 times with similar results. Dotted lines indicate the position of the peaks for the wild-type NF54/GDV1:GFP_cOE line. G) Quantification of the median fluorescence intensity of GDV1:GFP in induced or uninduced NF54/GDV1:GFP_cOE and NF54/GDV1Δ39:GFP_cOE parasite lines, normalised to uninfected parasites from each experiment, ± SD, n = 4. (*, p<0.05; ns, not significant, p≥0.05; NF54/GDV1:GFP_cOE uninduced vs induced, p=0.0227; NF54/GDV1Δ39:GFP_cOE uninduced vs induced, p=0.0318; NF54/GDV1:GFP_cOE induced vs NF54/GDV1Δ39:GFP_cOE induced p=0.0227; NF54/GDV1:GFP_cOE uninduced vs NF54/GDV1Δ39:GFP_cOE uninduced, p=0.1235); Statistical analysis was performed using Holm-Sidak corrected multiple comparison analysis of variance (ANOVA).

## Discussion

The aim of this study was to identify non-essential kinases as regulators of gametocyte commitment/development in *P. falciparum*. While several parasite lines of the kinase knock-out collection ^14^ were able to form gametocytes, two kinase KO lines showed a gametocytogenesis phenotype that led to the identification of a nonsense mutation in *gdv1* that results in a 39 aa truncation of the GDV1 C-terminus. This mutation may have been acquired by the common parental line prior to generation of the original transgenic lines, although several other clones from the Solyakov study we tested here are able to form gametocytes, possibly reflecting that only a proportion of the parasite population in the parental line carried the mutation. Alternatively, it cannot be excluded that the mutation arose independently in these two lines. This might be clarified by carrying out WGS, but this lies outside the scope of the present study. Our results show that the premature stop codon mutation in *gdv1* resulting in a 39 amino acid C-terminal truncation in the *tlk2* and *elk2* KO lines is sufficient to abolish sexual commitment. Based on our analysis of inducible GDV1 overexpression lines we observed that the truncated GDV1Δ39-GFP protein was present at substantially reduced protein levels compared to full-length GDV1-GFP. We propose that it is the loss of GDV1 stability that is the underlying cause for the lack of gametocytes in the GDV1 mutants. Although we have not further tested this here, the reduced amount of GDV1 protein shown in Western blots is likely caused by a destabilisation of the GDV1 protein due to the truncation. Alternatively, although unlikely, it could be caused by a reduction in *gdv1* transcripts. Regardless of the observed decrease of GDV1 protein levels, the truncation does neither result in a strong nuclear localisation defect when overexpressed as a GFP fusion protein, nor in a failure to interact with HP1 expressed in bacteria. It will be important to show in the future whether the few NF54/GDV1Δ39:GFP_cOE parasites, which show similar levels of GDV1Δ39:GFP compared to GDV1:GFP in NF54/GDV1:GFP_cOE parasites, are able to induce gametocytogenesis. If they fail to do so, it would indicate additional functions of the GDV1 C-terminus, potentially contributing to bringing GDV1 to the *ap2-g* locus. In this respect, it would be of great interest to identify possible interactors of the GDV1 c-terminus.

## Material and Methods

### Plasmid construction and transfection

The construction of each of the ePK knockout plasmids here characterized has been described in ^14^. The *pMK-RQ-tkl2-loxPint* donor plasmid (synthesized by Geneart) contains a recodonized version of sequence containing the glycine-rich loop in the kinase domain of *tkl2* (rc. Gly Loop) flanked by two loxPints and homology regions for homology-directed repair. The *pDC2-Cas9-hDHFRyFCU* guideRNA plasmid targeting *tkl2* locus (pDC2_TKL2_gRNA) was generated using the primer pairs pDC2_TKL2_gRNA1_FOR/pDC2_TKL2_gRNA1_REV. Because we didn’t have a 3D7::DiCre line, we generated the 3D7/TKL2:loxPint conditional KO line by doing, for the first time, a double transfection with the *pMK-RQ-tkl2-loxPint* and pDC2_TKL2_gRNA, together with the pBSPfs47DiCre (containing the DiCre cassette) and the CRISPR/Cas9 plasmid pDC287 containing the guide RNA targeting the Pfs47 locus, as previously described ^23^. The plasmids were suspended in 100uL of P3 primary cell solution, 40ug of each rescue plasmid and 20ug of *pDC2-Cas9-hDHFRyFCU guide RNA* for each respective rescue plasmid, and transfected into the 3D7 parasites. *Briefly,* purified *P. falciparum* 3D7 schizont stages were electroporated using using Amaxa 4D-Nucleofector™ (Lonza) - program FP158 ^33^. Selection of parasites transfected was done using 5nM WR99210 (Jacobus Pharmaceutical) and after a first round of selection, cloned.

To generate the *pMK-RQ-gdv1Δ39-HA* plasmid, which upon integration into the endogenous *gdv1* locus mimics the mutation found in the kinase KO lines, the *gdv1* (PF3D7_0935400) 3’ homology region was PCR amplified from NF54 genomic DNA with primers #268/#269 (Supplementary Table 2). The amplified PCR fragment was Gibson-cloned into an AfIIII-digested plasmid synthesized by Geneart that contains a *gdv1* 5’ homology sequence followed by a recodonized truncated *gdv1Δ39* version and the sequence encoding the 3xHA tag (Supplementary Table 2). To generate the pD_cg6_cam-gdv1Δ39-gfp-glmS plasmid we amplified the gdv1Δ39 sequence from the pMK-RQ-gdv1Δ39-HA plasmid using primers #383/#384 (Supplementary Table 2) and introduced the PCR fragment using Gibson assembly into the donor plasmid pD_cg6_cam-gdv1-gfp-glmS ^29^ digested with EagI and BsaBI. The guideRNA cassette to mutate endogenous gdv1 was generated using the primer pairs pDC2_GDV1Δ39_gRNA1_FOR/pDC2_GDV1Δ39_gRNA1_REV and cloned into the pDC2-Cas9-hDHFRyFCU plasmid as previously described ^22^. The rescue plasmid *pMK-RQ-gdv1Δ39-HA* and the CRISPR/Cas9 plasmid *pDC2-Cas9-hDHFRyFCU* were suspended in 100uL of P3 primary cell solution, 40ug and 20ug DNA respectively, and transfected using Amaxa 4D-Nucleofector™ (Lonza). Briefly, purified *P. falciparum* NF54::DiCre schizont stages were electroporated using program FP158 ^33^. Selection of parasites transfected was done using 5nM WR99210 (Jacobus Pharmaceutical) and after a first round of selection, cloned. Transfection of NF54 parasites using the CRISPR/Cas9 pHF_gC-cg6 suicide plasmid ^29^ and the *pD_cg6_cam-gdv1Δ39-gfp-glmS* donor construct was performed as described previously ^12^. 50 µg each of the suicide plasmid and donor plasmid were transfected and parasites cultured in the presence of glucosamine to block NF54/GDV1Δ39:GFP_cOE protein overexpression. 24 hours after transfection and for six subsequent days in total, the transfected populations were treated with 4 nM WR99210 and then cultured in absence of drug selection until a stably propagating transgenic population was obtained. All primers, guide RNAs and fragments used in the construction and integration of the constructs as well as confirmation of rapamycin mediated excision are described in Supplementary Table 2.

### *Plasmodium falciparum* in vitro culture of asexual and sexual blood stages

*Plasmodium falciparum* parasite lines used in this study were all derived from the NF54 strain (originally isolated from an imported malaria case in the Netherlands in the 1980s; BEI Resources, cat. no. MRA-1000) ^34^. Asexual parasites were cultured in human blood (UK National Blood Transfusion Service) and RPMI 1640 medium containing 0.5% w/v AlbumaxII (Invitrogen) at 37°C, as previously described ^35^. Asexual parasites were used to produce gametocytes by seeding asexual rings at 1% or 3% parasitaemia and 4% haematocrit on day 0 and feeding the parasites once a day during 15 days (day 0 to day 14) in 3% O_2_-5% CO_2_-92% N_2_ gas, in RPMI complemented with 25mM HEPES, 50mg/liter hypoxanthine, 2g/L sodium bicarbonate, 10% human serum ^35,36^.

### Plasmodium falciparum sexual induction

Sexual induction of parasite lines was done by following Trager protocol ^35^. More specifically, gametocyte induction was started with a 3% asexual ring culture where sexual commitment was induced by using 50% spent medium, expecting the sexually committed merozoites to invade and develop during the next cycle ^20,35^. The overexpressing NF54/GDV1:GFP_cOE and NF54/GDV1Δ39:GFP_cOE parasite lines were kept in the constant presence of 2.5 mM glucosamine to block ectopic GDV1 expression and therefore sexual induction, while sexual induction was achieved culturing the parasites in the absence of glucosamine, as previously described ^29^.

### Time course of gametocyte induction, RNA extraction and RNA-seq library preparation

The samples were collected during the asexual cycle at 28-32 hpi and in the matching cycle at 28-32 hpi after induction of sexual commitment. The infected RBCs pellets were collected at the respective time point, centrifuged and solubilized in ten volumes of TRIzol (Ambion) prewarmed to 37°C, lysed for 5 minutes by mixing vigorously at 37°C and immediately frozen at −80°C until extraction. Complete RNA was isolated from the samples using Trizol/chloroform extraction followed by isopropanol precipitation^22^ and its concentration and integrity was verified using Agilent Bioanalyzer (RNA 6000 Nano kit) and NanoDrop 1000 spectrophotometer. 1-2 μg of total RNA from each sample (or complete sample if the yield was lower) was used for mRNA isolation (Magnetic mRNA Isolation Kit, NEB). First strand cDNA synthesis was performed using the SuperScript III First-Strand Synthesis System and a 1:1 mix of Oligo(dT) and random primers (Invitrogen). The DNA-RNA hybrids were purified using Agencourt RNACleanXP beads (Beckman Coulter) and the second cDNA strand was synthetized using a 10 mM dUTP nucleotide mix, DNA Polymerase I (Invitrogen) and RNAseH (NEB) for 2.5 h at 16°C. The long cDNA fragments were purified and fragmented using a Covaris S220 system (duty cycles = 20, intensity = 5, cycles/burst = 200, time = 30s). The ~200 bp long fragments were end-repaired, dA-tailed and ligated to “PCR-free” adapters ^37^ with index tags using NEBNext according to the manufacturer’s instructions. Excess adapters were removed by two rounds of clean-up with 1 volume of Agencourt AMPure XP beads. Final libraries were eluted in 30 μl water, quality-controlled using Agilent Bioanalyzer (High Sensitivity DNA chip) digested with USER enzyme (NEB) and quantified by qPCR. For some libraries additional 5 cycles of PCR amplification were performed, using KAPA HiFi HotStart PCR mix and Illumina tag-specific primers to obtain enough material for sequencing. Pools of indexed libraries were sequenced using an Illumina HiSeq2500 system (100 bp paired-end reads) according to manufacturer’s manual. All samples were generated in duplicates or triplicates and uninduced controls were always generated and processed in parallel. Raw data is available through GEO database repository (study GSE158689).

### RNAseq data analysis

The generation of raw data in the form of *.cram files quality control and adapter trimming was performed using the default analysis pipelines of the Sanger Institute. The raw data was transformed into paired *. fastq files using Samtools software (ver. 1.3.1). The generated reads were re-aligned to *Plasmodium falciparum* genome (PlasmoDB-30 release) in a splice aware manner with HISAT2 ^38^ using --known-splicesite-infile option within the splicing sites file generated based on the current genome annotation. Resulting *.bam files were sorted and indexed using Samtools and inspected visually using Integrated Genome Viewer (ver. 2.3.91). HT-seq python library ^38^ was used to generate reads counts for all genes for further processing. Raw counts were normalised to median-ratio and then tested against linear models of time nested in line and line nested within time using a negative binomial model for the normalised counts using DESeq2, differential genes being selected for a false discovery rate of < 0.1 ^39^.

### Saponin lysis and whole cell, cytoplasmic and nuclear protein extraction

10mL parasite culture (2-5% parasitemia, 4% haematocrit) was transferred to a 15 mL tube and centrifuged at 600 g for 5 min. The supernatant was aspirated and the RBC pellet resuspended in 5 volumes 0.15% saponin solution (2.5 mL for 500 μL RBC). After an incubation on ice of max. 10 min, the parasites were centrifuged at 1503 g for 5 min at 4°C. Subsequent steps were performed on ice in order to prevent protein degradation. The supernatant was aspirated and the parasite pellet resuspended in 1 mL cold phosphate buffered saline (PBS) and transferred to an Eppendorf tube. The parasite pellet was centrifuged at 1503 g for 30 sec at 4°C and washed with cold PBS until the supernatant was clear.

For whole cell protein extraction, one pellet volume (30-50 μL) of whole cell protein lysis buffer (8 M Urea, 5% SDS, 50 mM Bis-Tris, 2 mM EDTA, 25 mM HCl, pH 6.5) complemented with 1x protease inhibitor cocktail (Merck) and 1 mM DTT was added to the pellet at RT in order to lyse the parasites. The tube was vortexed, heated to 94°C for 5 min, sonicated for 2 min (5 cycles of 30 sec ON/ 30sec OFF), vortexed and heated again. Subsequently, the protein sample was centrifuged at 20238 g for 5 min at RT and the supernatant was transferred into a new tube, which was frozen at −20°C and stored until use.

For cytoplasmic and nuclear protein extraction, the parasite pellet was lysed in 300μL cytoplasmic lysis buffer (20 mM Hepes (pH 7.9), 10 mM KCl, 1 mM EDTA, 0.65% Igepal) complemented with 1x protease inhibitor cocktail (Merck) and 1 mM DTT (leaving the nucleus intact) and incubated on ice for 5 min (Voss et al., JBC, 2002). The lysed parasites were centrifuged at 845 g for 3 min, the supernatant representing the cytoplasmic protein fraction was transferred into a new tube and placed on ice. The remaining nuclear pellet was washed in 500 μL cytoplasmic lysis buffer and centrifuged at 845 g for 3 min. The washing was repeated until the supernatant was clear. The nuclear pellet was resuspended in 60 μL whole cell lysis buffer and vortexed at high speed at RT for 10-20 min. The insoluble material was centrifuged at 20238 g for 3 min, the supernatant representing the nuclear protein fraction was transferred to a new tube and placed on ice. Both protein fractions were frozen at −20°C and stored until use.

### Western Blot

Parasite extracts were solubilized in protein loading buffer, denatured at 95 °C for 10 min, subjected to SDS-PAGE and transferred onto a nitrocellulose membrane. Membranes were immunostained with mouse anti-GFP (1:250 dilution; Roche, 11814460001), rabbit anti-Aldolase-HRP conjugated (1:5000 dilution, abcam ab38905) and rabbit anti-Histone 3 (1:2000 dilution, abcam ab1791) primary antibodies. Antibody detection was done using chemiluminescent western blot detection using goat anti-mouse secondary antibody conjugated with HRP and the ECL western blotting detection reagents (Amersham RPN2106) or by direct infrared fluorescence detection on the Odyssey Infrared Imaging System (Odyssey CLx, LI-COR) using IRDye 680LT goat anti-rat IgG (1:10000 dilution; LI-COR) and IRDye 800CW goat anti-rabbit IgG (1:10000 dilution; LI-COR).

### Immunofluorescence assay at different parasite stages

Air-dried thin blood films of asexual parasites were fixed with 4% paraformaldehyde containing 0.0075% glutaraldehyde for 15 min and permeabilized in 0.1% (v/v) Triton X-100 (Sigma) for 10 min ^40^. Blocking was per formed in 3% BSA for 1 h. Slides were incubated with rat anti-HA high-affinity (1:1000 dilution; Roche, clone 3F10) at room temperature for 30 min, followed by Alexa fluor conjugated goat anti-rat IgG (1:1000 dilution; Thermo Fisher Scientific) at room temperature for 30 min. Parasite nuclei were stained with 4’, 6-diamidino-2-phenylindole (DAPI; Invitrogen). Slides were mounted in ProLong^®^ Gold antifade reagent (Invitrogen) and images were obtained with the inverted fluorescent microscope (Ti-E; Nikon, Japan) and processed using NIS-Elements software (Nikon, Japan).

### Flow Cytometry

NF54/GDV1:GFP_cOE and NF54/GDV1Δ39:GFP_cOE parasites were grown in the presence or absence of glucosamine in order to block or allow sexual commitment, respectively. Schizonts from the 4 conditions were purified by percoll gradient and allowed to invade fresh red blood cells for 4h, before uninvaded schizonts were removed. Flow cytometry analysis was performed at approximately 44h post invasion, in 4 biological replicates. For one replicate, parasites were fixed for 1h in 4% paraformaldehyde in PBS, stained with Hoechst 33342 (1:1000 in PBS) for 10 minutes and analysed on an LSRFortessa flow cytometer (Becton Dickinson) using FACSDiva software. For the other three replicates, live parasites were stained with Hoeschst 33342 and analysed on a BD FACSAria II flow cytometer (Becton Dickinson) using FACSDiva software. Hoechst fluorescence was detected using a 355nm (UV) excitation laser with a 450/50nm bandpass filter, while GFP fluorescence was detected with a 488nm (blue) excitation laser, a 505nm longpass filter and a 530/30nm bandpass filter. At least 30,000 cells were counted for each sample. Data were analysed using FCS Express 7 (Research Edition) software. The population was first gated on single cells based on the side and forward scatter, then on highly Hoechst-positive infected schizonts, before the median fluorescence intensity (MFI) of the GFP-fluorescence was calculated for each line. An example of the gating strategy for infected cells is shown in Supplementary Figure 4. Due to the variation in fluorescence intensity between different experiments, MFI values were normalised by dividing the MFI of each infected sample by the average MFI of the uninfected samples within the same experiment (n=4). Statistical analysis was performed using Holm-Sidak corrected multiple comparison analysis of variance (ANOVA) on samples paired within each experiment using Graphpad Prism version 8.

### In vitro protein-protein interaction experiments

In order to co-express Strep(II)-tagged HP1 with a His-SUMO-tagged truncated version of GDV1, we deleted the 39 C-terminal amino acids of the coding sequence of GDV1 in the vector pStrep-HP1_HS-GDV1 ^12^. For this purpose, we circularized a PCR product amplified from this vector with the primers D39F and D39R using Gibson assembly. The proteins were expressed and the in vitro interaction assay was performed as previously described (14) using full-length GDV1 as positive and SIP2 as negative control.

## ACKNOWLEDGEMENTS

We would like to thank Christian Doerig for the original *Plasmodium falciparum* kinase KO lines. We would like to thank Frank Schwach and Mandy Sanders for preparing, running and initial quality control of RNAseq samples. We would like to thank Ellen Knuepfer and Christiaan van Ooij for the pBSPfs47DiCre and pDC287 plasmids, as well as scientific advice. We would like to thank Kostas Kousis for his help with the Flow Cytometry data acquisition.

## AUTHOR CONTRIBUTIONS

M. Tibúrcio and M. Treeck conceived the study. M. Tibúrcio performed most of the parasite genetic manipulations and all the parasite line phenotyping experiments, as well as RNAseq material collection. E. Hitz generated the NF54/GDV1Δ39:GFP_cOE parasite line, I. Niederwieser performed the in vitro protein-protein interaction experiments and T. S. Voss supervised these experiments and provided conceptual advice and resources. Gavin Kelly performed RNAseq analysis. RNAseq samples were run in Oliver Billkers group at the Sanger Institute. H. Davies performed the Flow Cytometry Data Analysis. Christian Doerig provided the original *P. falciparum* kinase KO cell lines. All authors contributed to experimental design and interpretation of the results. M. Tibúrcio and M. Treeck wrote the article with contributions from all authors.

---

**Supplementary Figure 1.**
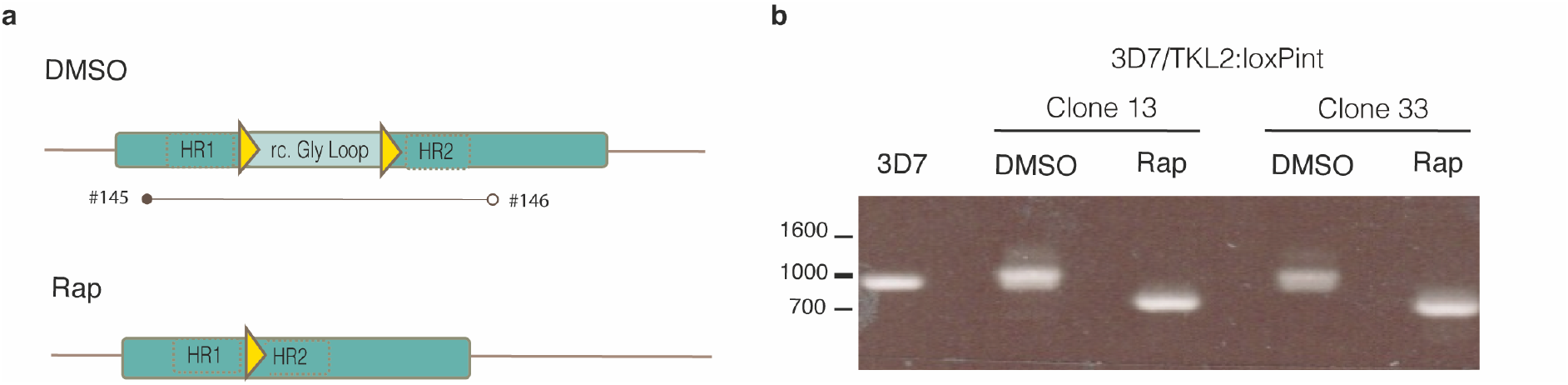
Confirmation of *tkl2-loxPint* cassette integration into the *Plasmodium falciparum* 3D7 line and the efficiency of DiCre mediated excision. A) Representation of the primer pairs used to test correct integration of *tkl2-loxPint* cassette and efficient rapamycin mediated excision. B) PCR analysis shows correct integration of *tkl2-loxPint* cassette and near complete excision of TKL2:LoxPint in two different clones from the same transfection (clone 13 and 33) after rapamycin treatment. The sequences of the primers used are in Supplementary Table 2.

**Supplementary Figure 2.**
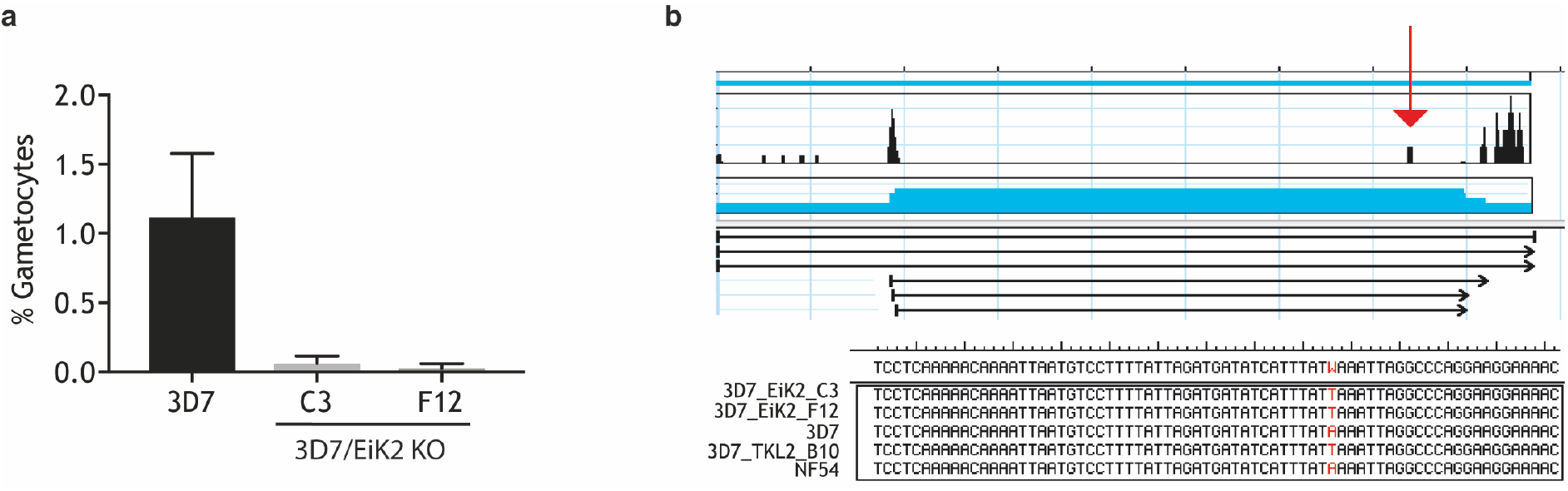
PfeIK2 kinase KO parasites fail to produce gametocytes. A) Comparison of gametocytemia upon sexual induction between the 3D7 WT line (n=4) and two PfeIK2 kinase KO clones (n=2 for each clone, both from the same transfection). Parasite line and clones measured in at least two biological experiments with single replicates. Each column represents the mean number of gametocytes in at least 500 cells. B) Representation of GDV1 sequences of the NF54, 3D7, PfeiK2 and TKL2 parasite lines and the identified point mutation in the PfeIK2 and PfTKL2 KO clones (red arrow) which is absent in the 3D7 and NF54 reference parasite lines.

**Supplementary Figure 3.**
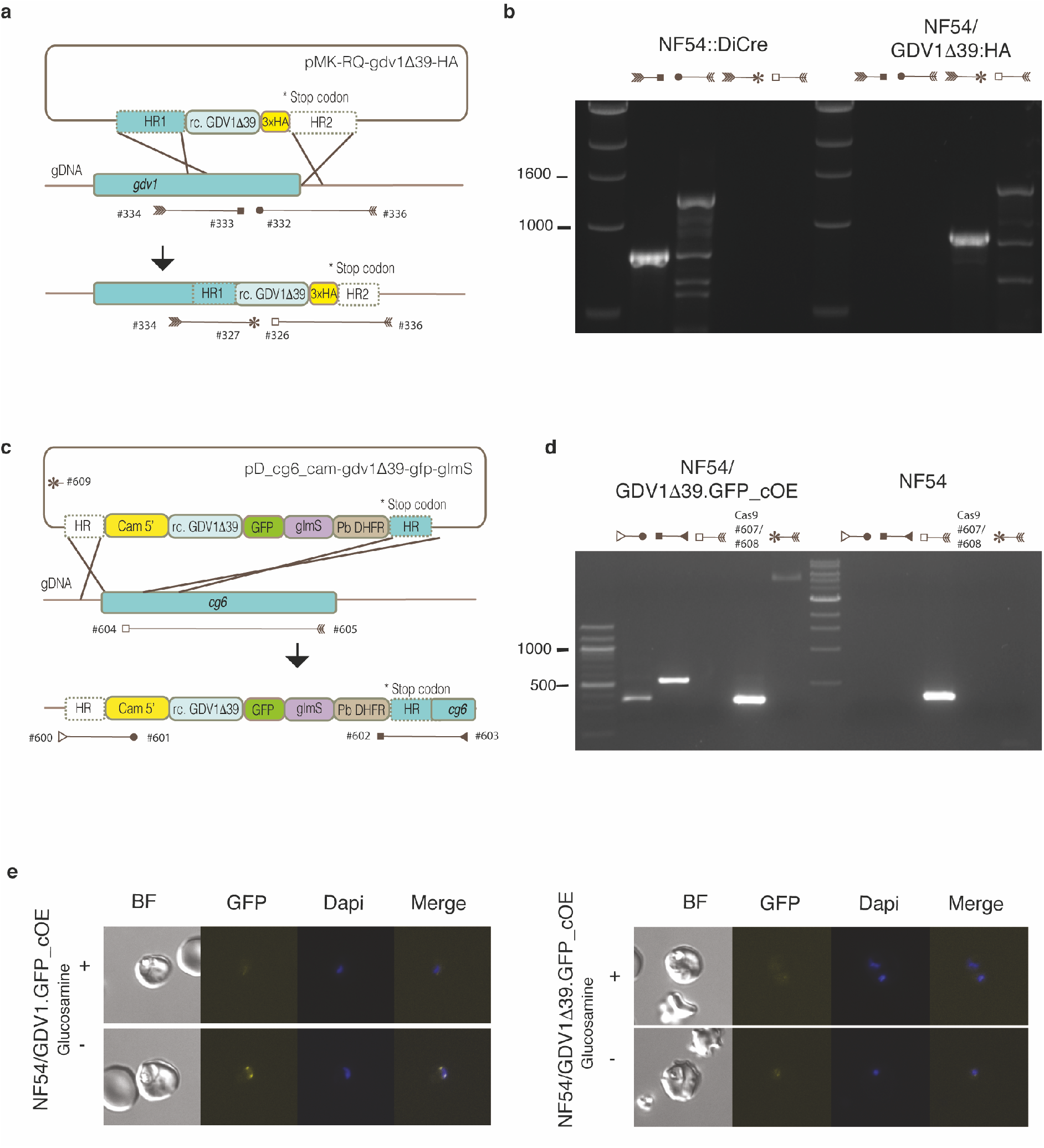
Generation of the NF54/GDV1β39:HA and NF54/GDV1Δβ39:GFP_cOE *Plasmodium falciparum* lines. A) Illustration of the strategy used to generate the 3xHA-tagged GDV1Δ39 mutant line (NF54/GDV1Δ39:HA) as well as the primers used to confirm donor sequence integration. B) PCR analysis of integration of the gdv1Δ39:ha construct into the *gdv1* locus in the NF54 *P. falciparum* parasite line. C) Schematic of the strategy used to generate the NF54/GDV1Δ39:GFP_cOE overexpressing line as well as the primers used to verify integration of the transgene cassette into the *cg6 (glp3*) locus. The pGDV1Δ39:GFP_cOE donor plasmid contains a 5’ Cam sequence followed by a recodonized version of the *gdv1Δ39* mutant in-frame with *gfp* sequence, the *glmS* ribozyme element and *Plasmodium berghei* dihydrofolate reductase (PfDHFR), flanked by two homology regions. D) PCR analysis of *gdv1Δ39-gfp-glmS* cassette integration in the cg6 locus in the NF54 *P. falciparum* parasite line and the presence of the CRISPR/Cas9/Suicide plasmid. E) Quantification of GFP expression in the GDV1:GFP and GDV1Δ39:GFP lines cultured in in the presence (prevents GDV1 expression) or absence of glucosamine (induces GDV1 overexpression).

**Supplementary Figure 4.**
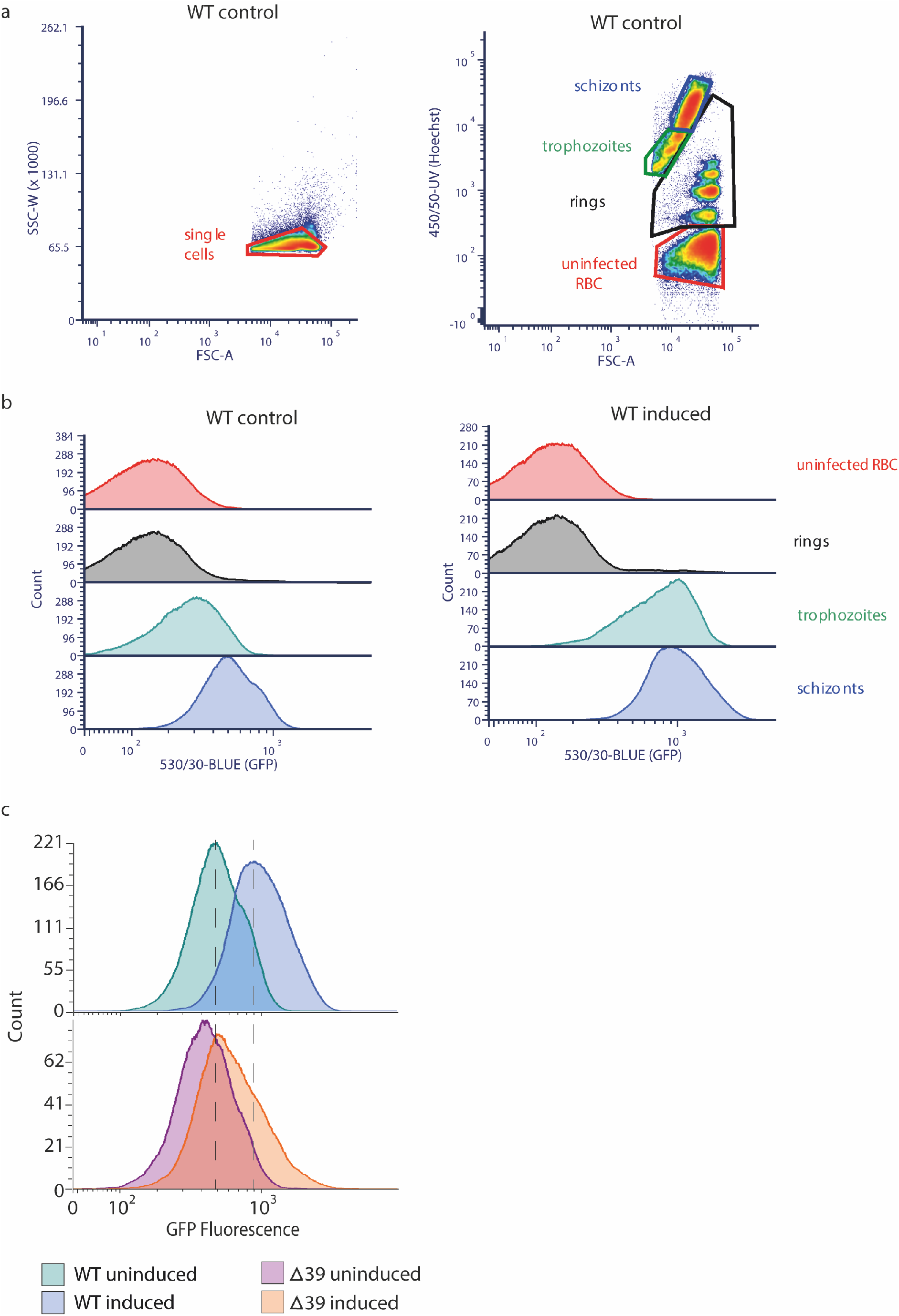
A) Gating strategy for flow cytometry experiments. B) Flow cytometry histograms quantifying GFP fluorescence in uninfected RBC and RBC infected with ring stage parasites, trophozoites and schizonts. Stacked plots are shown for both GDV1:GFP uninduced and induced parasites. C) Normalised flow cytometry histograms quantifying GFP fluorescence for each line. The experiment was repeated 4 times with similar results. Dotted lines indicate the position of the peaks for the wild-type NF54/GDV1:GFP_cOE line.

**Supplementary Table 1.** Comparison of gene expression, based on RNAseq data, of samples from 3D7 WT and PfeIK2 C3 and F12 kinase KO clones. Log2 fold changes of induced vs. non-induced parental (3D7) and the 2 *gdv1* mutant parasite clones from the PfeIK2 KO clones C3 and F12.

**Supplementary Table 2.**
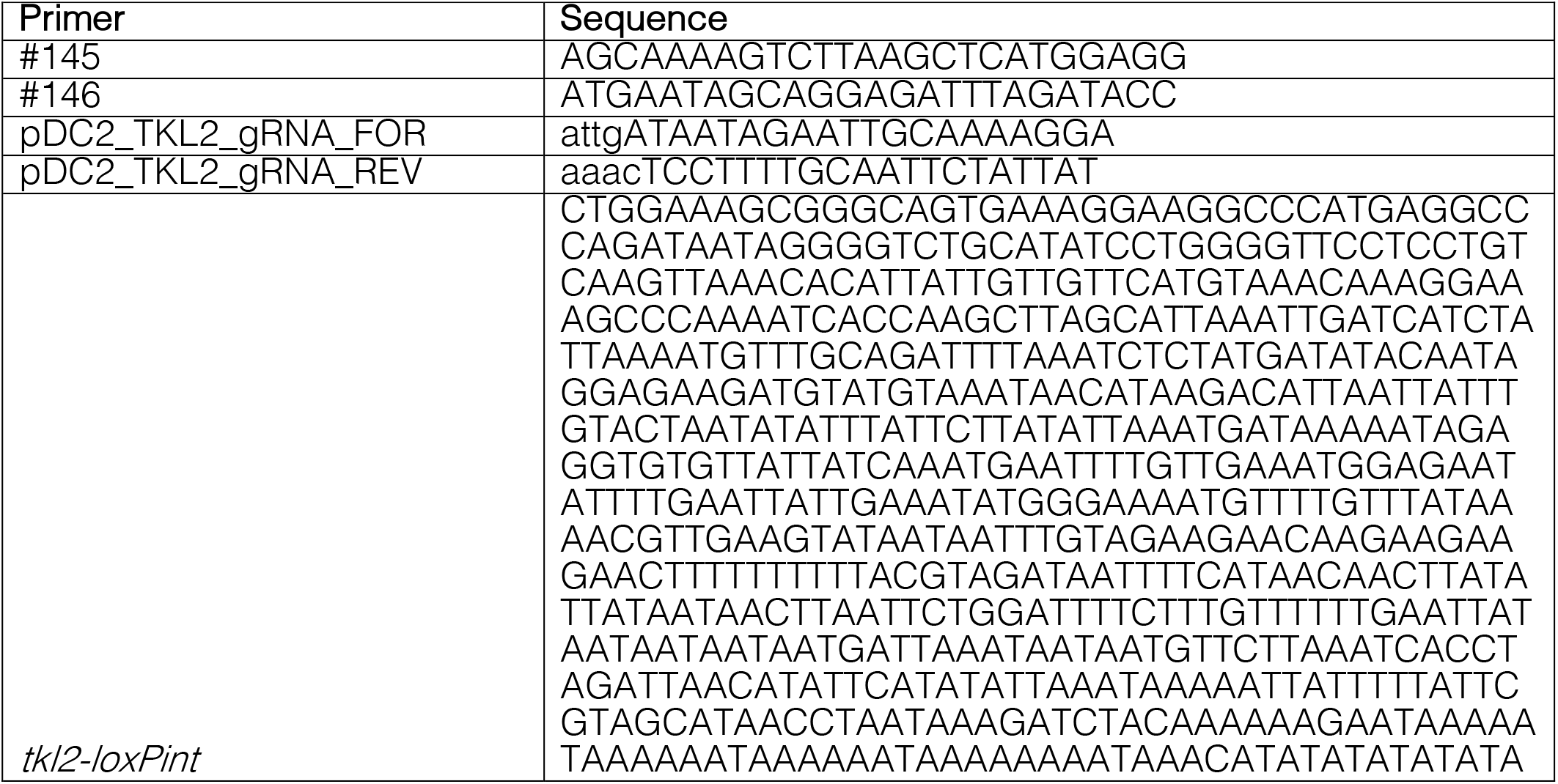
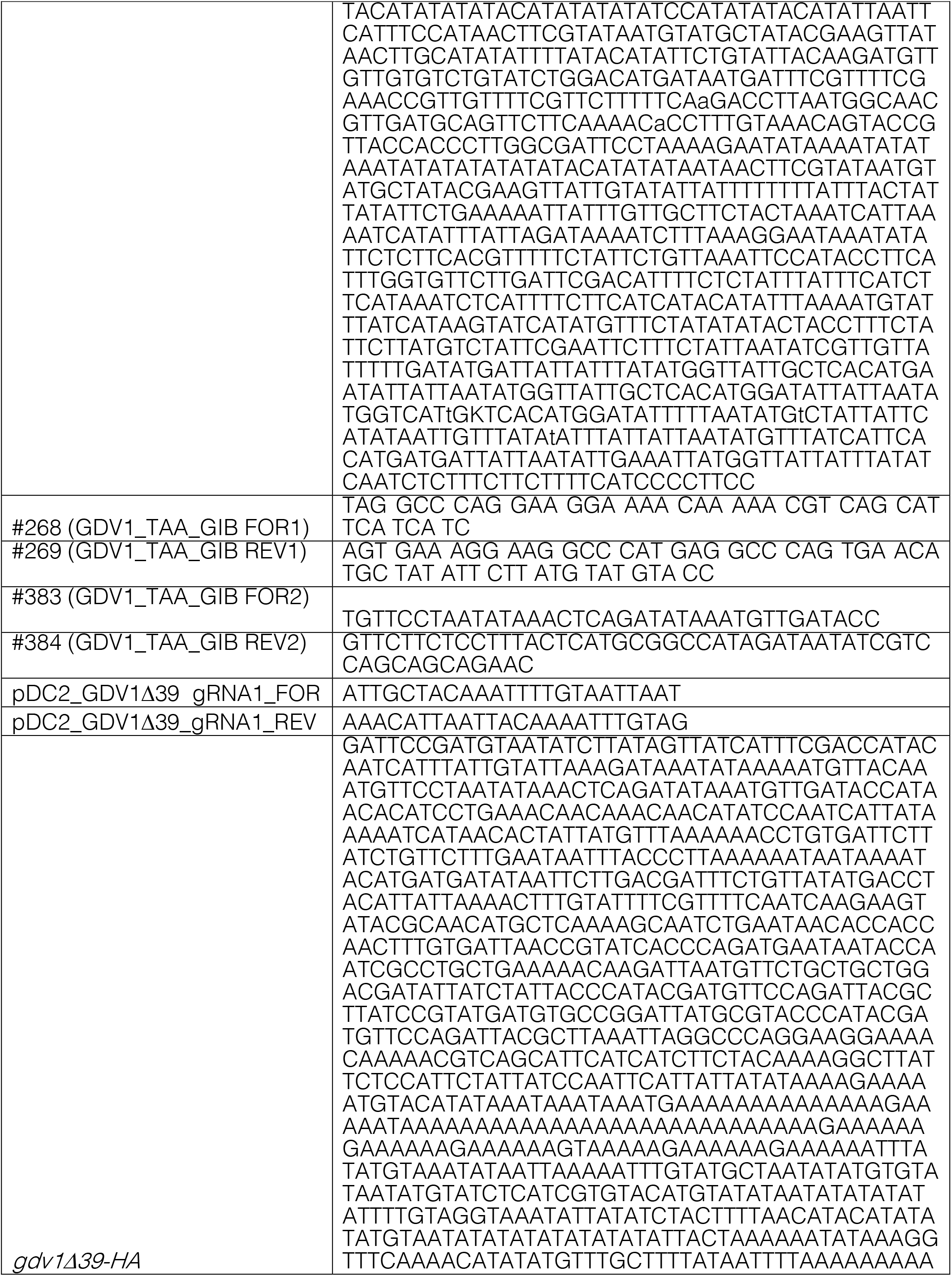
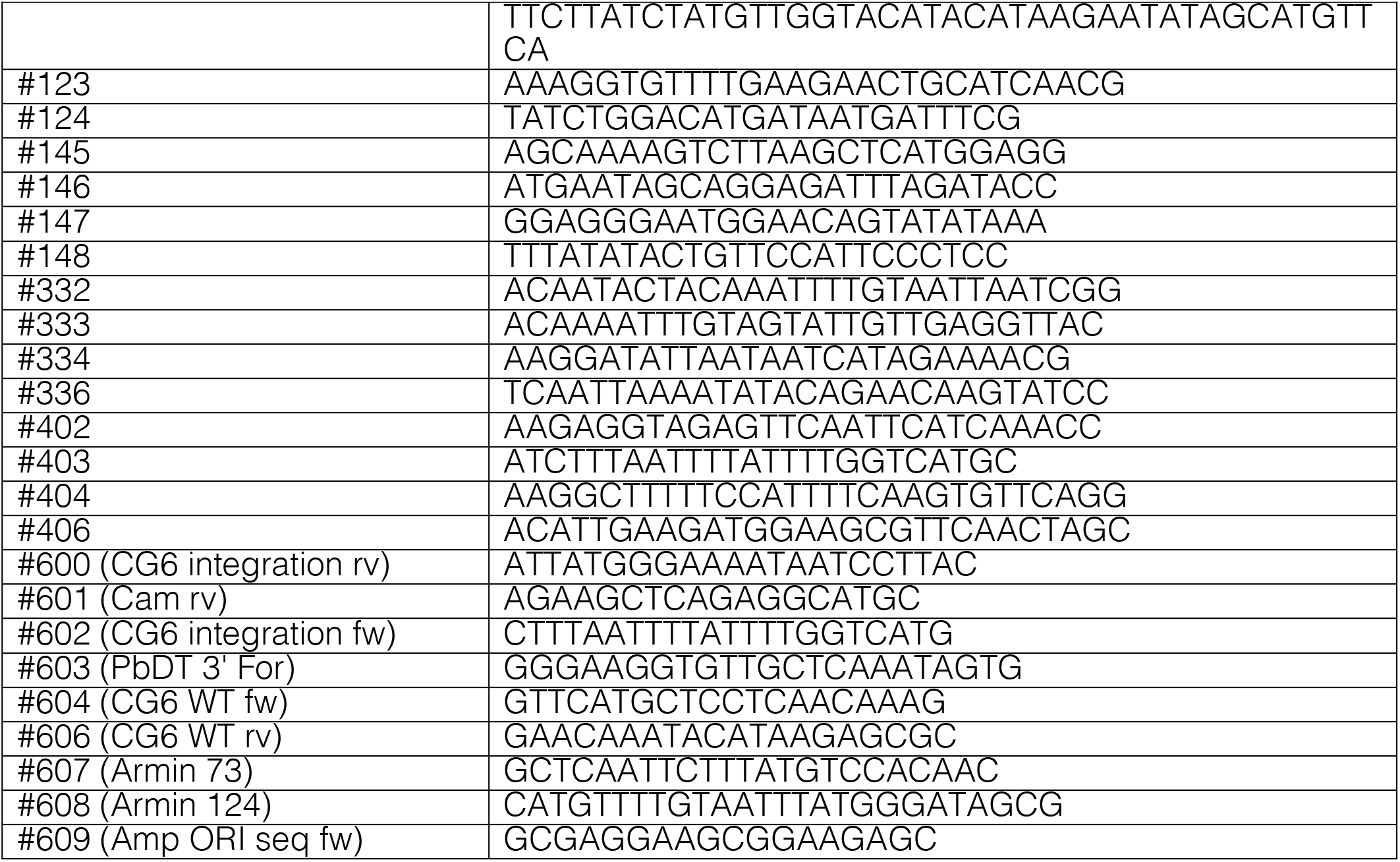
Primers, fragment and guide RNA sequences used to generate and confirm integration and rapamycin-induced excision of *pMK-RQ-tkl2-loxPint*, *pMK-RQ-gdv1Δ39-HA* and *pD_cg6_cam-gdv1**Δ**39-gfp-glmS*.

